# Emerging SARS-CoV-2 diversity revealed by rapid whole genome sequence typing

**DOI:** 10.1101/2020.12.28.424582

**Authors:** Ahmed M. Moustafa, Paul J. Planet

## Abstract

**Background:** Discrete classification of SARS-CoV-2 viral genotypes can identify emerging strains and detect geographic spread, viral diversity, and transmission events.

**Methods:** We developed a tool (GNUVID) that integrates whole genome multilocus sequence typing and a supervised machine learning random forest-based classifier. We used GNUVID to assign sequence type (ST) profiles to each of 69,686 SARS-CoV-2 complete, high-quality genomes available from GISAID as of October 20^th^ 2020. STs were then clustered into clonal complexes (CCs), and then used to train a machine learning classifier. We used this tool to detect potential introduction and exportation events, and to estimate effective viral diversity across locations and over time in 16 US states.

**Results:** GNUVID is a scalable tool for viral genotype classification (available at https://github.com/ahmedmagds/GNUVID) that can be used to quickly process tens of thousands of genomes. Our genotyping ST/CC analysis uncovered dynamic local changes in ST/CC prevalence and diversity with multiple replacement events in different states. We detected an average of 20.6 putative introductions and 7.5 exportations for each state. Effective viral diversity dropped in all states as shelter-in-place travel-restrictions went into effect and increased as restrictions were lifted. Interestingly, our analysis showed correlation between effective diversity and the date that state-wide mask mandates were imposed.

**Conclusions:** Our classification tool uncovered multiple introduction and exportation events, as well as waves of expansion and replacement of SARS-CoV-2 genotypes in different states. Combined with future genomic sampling the GNUVID system could be used to track circulating viral diversity and identify emerging clones and hotspots.

## Introduction

Rapid sequencing of the SARS-CoV-2 pandemic virus has presented an unprecedented opportunity to track the evolution of the virus and to understand the emergence of a new pathogen in near-real time. During its explosive radiation and global spread, the virus has accumulated enough genomic diversity that we are now able to identify distinct lineages and track their spread in distinct geographic locations and over time (Bedford, et al. 2020; Chen, et al. 2020; Deng, et al. 2020; Rambaut, et al. 2020; Shen, et al. 2020; Worobey, et al. 2020). Phylogenetic analyses in combination with rapidly growing databases (Shu and McCauley 2017; Rambaut, et al. 2020) have been instrumental in identifying distinct clades and tracing how they have spread across the globe, as well as estimating calendar dates for the emergence of certain clades (Bedford, et al. 2020; Deng, et al. 2020; Rambaut, et al. 2020; Worobey, et al. 2020). This information is extremely useful in assessing the impact of early measures to combat spread as well as identifying missed opportunities (Korber, et al. 2020; Worobey, et al. 2020).

Although reconstructing a robust phylogeny of viral variants is an intuitive approach for viral classification, traditional phylogenetic approaches suffer from problems with scalability. Building comprehensive phylogenetic trees for single nucleotide polymorphism (SNP) based analysis of SARS-CoV-2 is already extremely computationally expensive, and will become more and more difficult as hundreds of thousands of sequences are added. Dividing the dataset into subsets of genomes necessarily loses information and explanatory power. Because of this roadblock, our goal was to develop a rapid way to categorize genomes that scales readily and leads to as little information loss as possible. We saw an opportunity to combine our allele identifying tool, WhatsGNU (Moustafa and Planet 2020b), with the Multilocus Sequence Typing (MLST) approach (Maiden, et al. 1998) that has been widely used in bacterial classification, tracking the emergence of new lineages, and associating specific Sequence Types/Clonal Complexes (STs/CCs) with certain diseases. Our whole genome MLST (wgMLST) approach rapidly assigns an allele number to each gene nucleotide sequence in the virus’s genome creating a sequence type (ST), which is codified as the sequence of allele numbers for each of the ten genes in the viral genome.

Here we show that this approach allows us to link STs into clearly defined clonal complexes (CC) that are consistent with phylogeny and other SARS-CoV-2 typing systems (Shu and McCauley 2017; Rambaut, et al. 2020). We show that assessment of STs and CCs agrees with multiple introductions of the virus in certain US states. In addition, we use temporal assessment of ST/CC diversity to uncover waves of expansion and decline, and the apparent replacement of certain CCs with emerging lineages in specific US states.

## Results and Discussion

We developed the GNU-based Virus IDentification (GNUVID) system as a tool that automatically assigns a number to each unique allele of the ten open reading frames (ORFs) of SARS-CoV-2 (Wu, et al. 2020) (Figure 1A). GNUVID compressed the 696,860 ORFs in 69,686 high quality GISAID genomes (Supplementary Table 1) to 37,921 unique alleles in five minutes on a standard desktop, achieving 18-fold compression and losing no information. To create an ST for each isolate GNUVID automatically assigned 35,010 unique ST numbers based on their allelic profile (Supplementary Table 1). We then used a minimum spanning tree (MST) to group STs into larger taxonomic units, clonal complexes (CCs), which we define here as clusters of >20 STs that are single or double allele variants away from a “founder”. Using the goeBURST algorithm (Feil, et al. 2004; Francisco, et al. 2009) to build the MST and identify founders, we found 154 CCs (Figure 1A and Supplementary Table 1).

**Figure 1.**
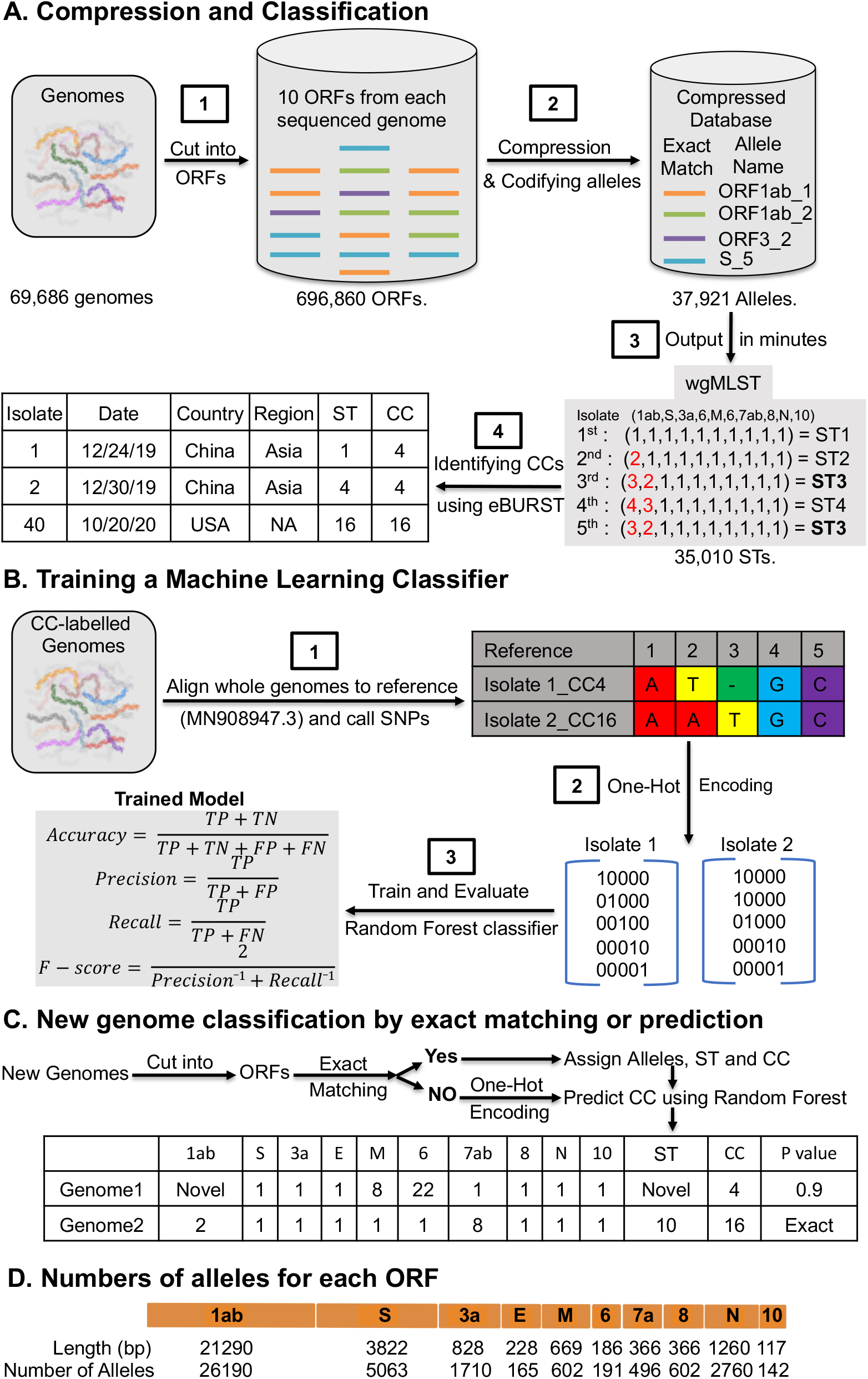
Workflow for the GNUVID tool and its compression technique. **A. Compression and classification**. The tool starts by compressing the database of the 10 ORFs of each of the SARS-CoV-2 genomes to only include a unique sequence for each allele type. The tool then uses a whole genome multilocus sequence typing (wgMLST) approach by assigning an allele number to each gene nucleotide sequence in the virus’s genome creating a sequence type (ST) which is codified as the sequence of allele numbers for each of the ten genes in the viral genome. The STs are then linked into clearly defined clonal complexes (CCs) using goeBURST. **B. Training a machine learning classifier**. The CC-labelled genomes are then aligned to the SARS-CoV-2 reference genome (MN908947.3) and single nucleotide polymorphisms (SNPs) are called. The SNP matrix is then one-hot encoded and used to train a random forest classifier. The training followed a 5-fold cross-validation approach to assess the prediction capabilities of GNUVID according to four statistics (accuracy, precision, recall and F-score). TP, TN, FP and FN are true positives, true negatives, false positives and false negatives, respectively. **C. New Genome classification by exact matching or prediction.** GNUVID first tries to match each of the 10 ORFs from a query SARS-CoV-2 genome to an exact match in the compressed database to define an ST, and matches that to any associated CC. If no exact match is found due to novelty or ambiguity in any of the 10 ORFs, the query genome is aligned to the reference, one-hot encoded and a CC is predicted by the trained classifier. A report is then created showing the allele number of each ORF, ST, CC and a probability of membership in the CC. **D.** Map of SARS-CoV-2 virus genome showing the length in base pairs (bp) of the ten ORFs and numbers of alleles in the current database 69,686 isolates. The majority of the identified 37,921 unique alleles (69%) are for ORF1ab which represents 71% of the genome length. Strikingly, the two highest ratios (number of alleles/ORF length) are for the nucleocapsid protein (2.2) and ORF3a (2.1) while the spike protein had a ratio of 1.32.

A random forest classifier was then trained on 53,565 CC-labelled genomes. The overall prediction statistics of the model were accuracy: 0.955, F-score: 0.950, precision: 0.947, and recall: 0.964 (Figure 1B).

For any new query genome, GNUVID attempts to classify it first by exact matching of the allelic profile to one of the other STs. If there is no exact match, the CC for the query genome is predicted using the trained model. This query process saves time and also allows each ORF to be typed and tallied individually (Figure 1C and 1D).

To show that CCs are mostly consistent with whole genome phylogenetic trees, we mapped the 10 most common CC designations onto a maximum likelihood tree. Members of the same CC usually grouped together in clades (Supplementary Figure 1). To further validate our wgMLST classification system we compared it to the proposed “dynamic lineages nomenclature” for SARS-CoV-2 (Rambaut, et al. 2020) and GISAID clades naming system (Shu and McCauley 2017). A high percentage of CCs, 95.5% (147/154) and 87.7% (135/154) of the CCs, had 90% of their genomes assigned to the same GISAID clade and pangolin lineage, respectively, showing strong agreement between these classification schemes (Supplementary Table 1). One limitation of our classification strategy, as with many schemes that operate in real time, is that paraphyletic groups can occur as a new ST arises from an older ST (e.g. CC258 and CC768 emerged from CC255 and CC258 making CC255 and CC258 paraphyletic, respectively) (Supplementary Figure 1). While this means that not all ST/CC groups will be monophyletic, this property of the nomenclature may be helpful in gauging emergence and replacement of an ancestral form.

When the global region of origin for each genome sequence was mapped to each CC there was a strong association of later emerging CCs with certain geographical locations, possibly reflecting relative containment after international travel restrictions (Figure 2). To obtain an up-to-date picture of virus diversity in the US, we analyzed 107,414 high coverage genomes (isolation dates between December 2019 to October 20^th^ 2020) from the GISAID (Supplementary table 1). There were 26,528 genomes isolated in the US in this dataset that belong to 87 of 154 CCs. Strikingly, 35% of the genomes belong to CC258 (GISAID clade GH) and 75% of the genomes are represented by just 10 CCs (CC4, 255, 256, 258, 300, 498 768, 3530, 10221, 21210)). Moreover, 72% (63/87) of the CCs (representing 82% of the genomes) had the spike D614G mutation that has been associated with increased spread (Korber, et al. 2020). Interestingly, none of the US genomes were associated with any of the 12 CCs (26377, 26754, 27693, 27950, 28012, 28825, 29259, 29310, 30362, 31179, 31744 and 31942) that have the spike protein A222V mutation (GISAID clade GV) that has been recently associated with increased spread in the Europe (Hodcroft, et al. 2020). Ten of the 12 CCs with the A222V mutation were isolated only from Europe while the two other CCs (27693 and 27950) had 2 genomes from Hong Kong and 6 from New Zealand, respectively. This shows a strong association of this clade with Europe.

**Figure 2.**
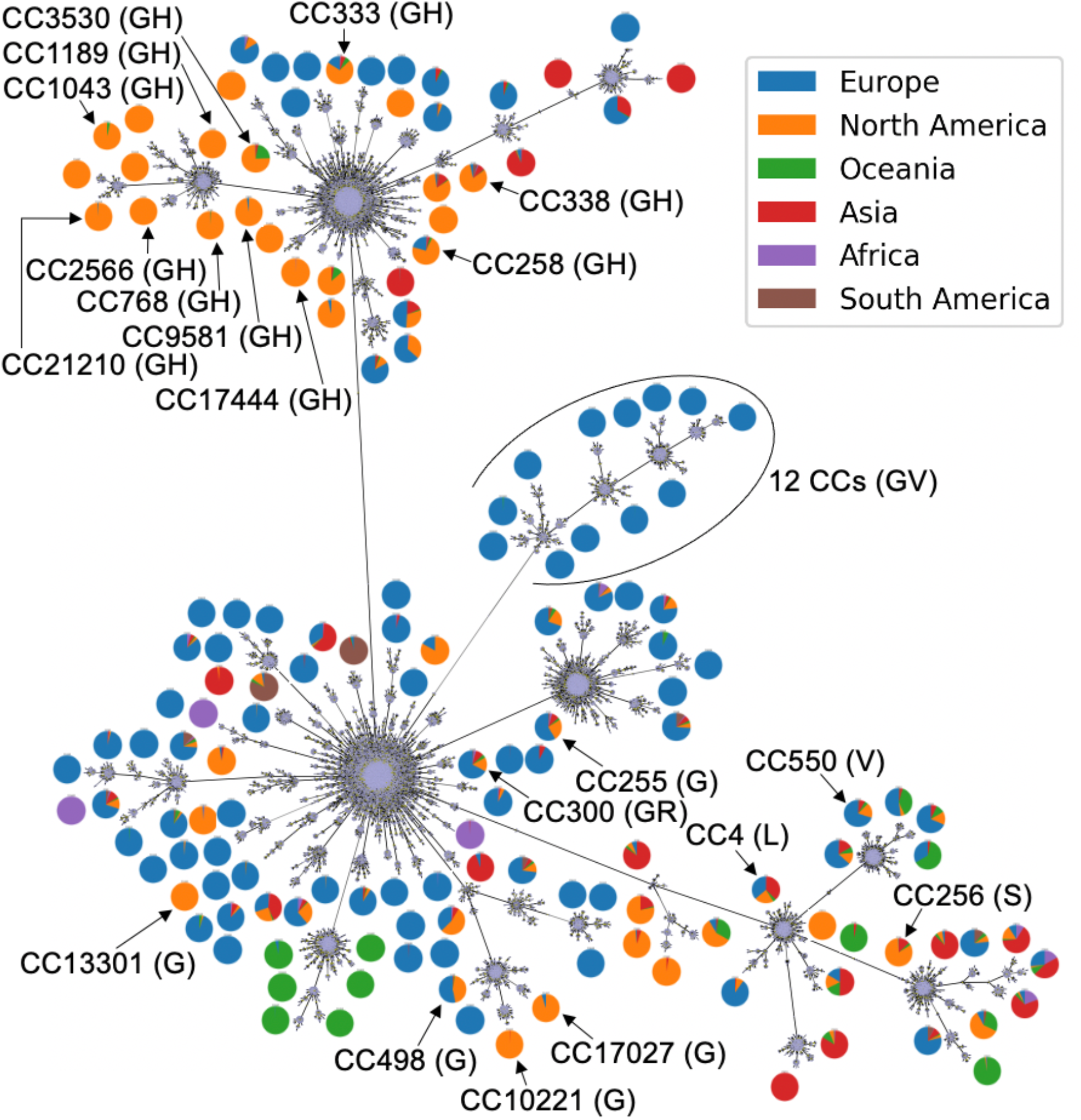
Global SARS-CoV-2 Diversity. Minimum spanning tree from goeBURST of the 35,010 Sequence Types (STs) showing the 154 Clonal Complexes (CCs) identified in the dataset. Only the most common 20 CCs in the 16 states are shown in black. The pie charts show the percentage of genomes from the different geographic regions in each CC.

The relative proportions of STs or CCs isolated and sequenced may be a highly biased statistic that is contingent upon where the isolate comes from, the decision to sequence its genome, and the local capacity to sequence a whole genome. Certain states (Washington, Texas and California) clearly sequenced more genomes than the other states. Focusing on specific states may help to partially ameliorate this bias, and we chose to focus on 16 states (Washington (WA), Texas (TX), California (CA), Wisconsin (WI), New York (NY), Michigan (MI), Minnesota (MN), Louisiana (LA), Utah (UT), Virginia (VA), Florida (FL), Oregon (OR), Massachusetts (MA), New Mexico (NM), Maryland (MD), and Connecticut (CT)) with at least 200 genomes in the studied time period, representing 92.6% (24,565/26,528) of all viral genomes available from the US. The most common 20 CCs in these states, representing 86.5% (21261/24565) of the genomes, are shown in Figure 2.

Because we included collection dates for each genomic sequence, we can use STs and CCs to better understand the emergence and replacement of certain lineages and viral diversity in geographical regions over time. Figure 3A and Supplementary Figure 2 show temporal plots of the most common 20 CCs in 16 states. In WA, the earlier introduction CC256 (GISAID clade S) was replaced by CC258 (GISAID clade GH), perhaps by introduction from the East Coast or Europe (Bedford, et al. 2020; Deng, et al. 2020). CC258 was then replaced by CC300 (GISAID clade GR) and subsequently by CC498 (GISAID clade G).

**Figure 3.**
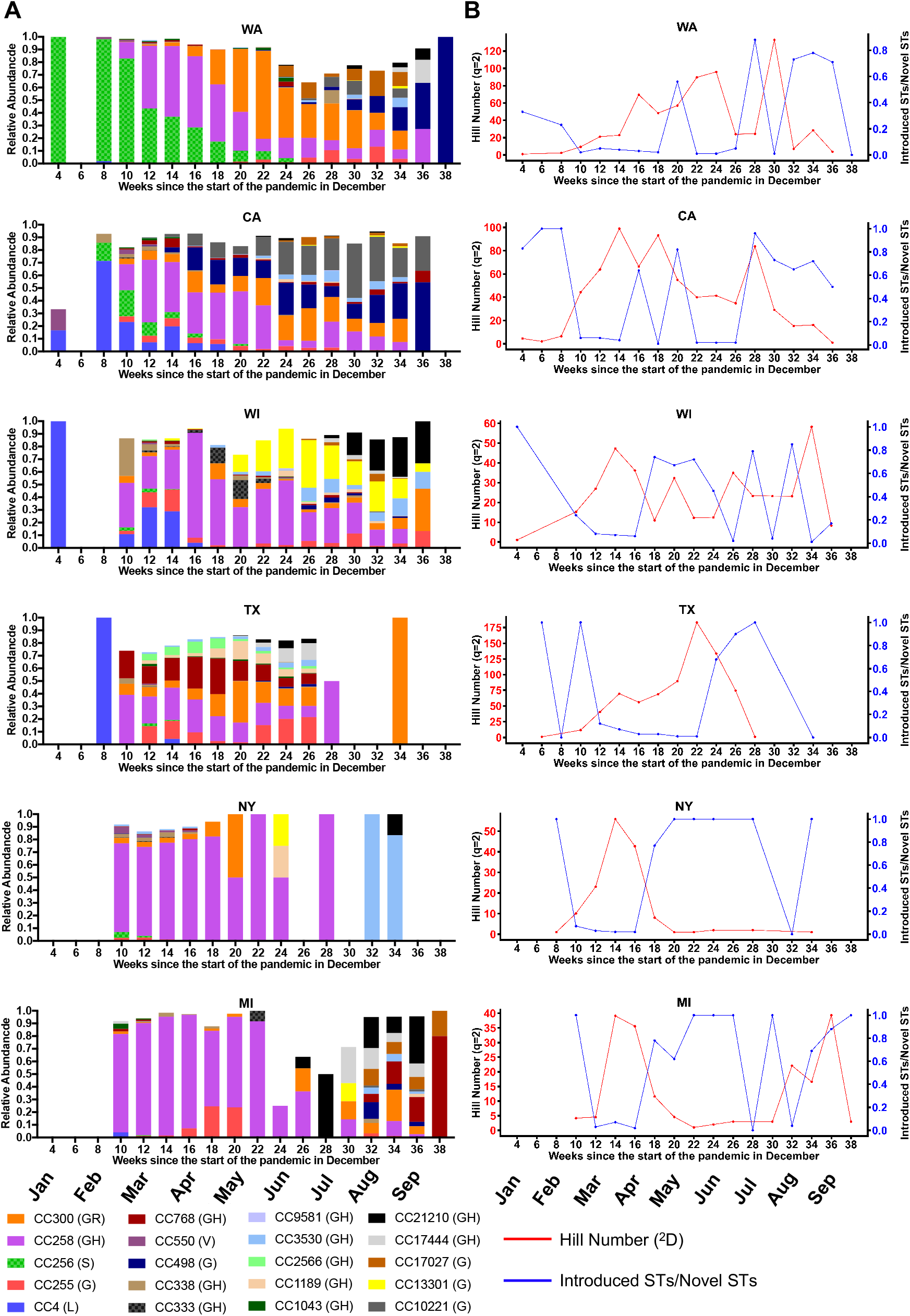
SARS-CoV-2 diversity in 6 states over time. **A.** Temporal Plots of circulating Clonal Complexes and corresponding GISAID clade in parentheses in six different states (Washington (WA), California (CA), Wisconsin (WI), Texas (TX), New York (NY) and Michigan (MI)). The visualizations were limited to the 20 most common CCs. **B.** Diversity of Sequence Types (STs) in the six states over time are represented for each 2-week time period in the following ratios: 1. Effective diversity (Hill number equivalent (^2^D) of Simpson index (^2^H)) (red) 2. Number of STs new to a state that were previously isolated and sequenced outside a state divided by the number of STs not seen previously in a state (blue).

In the neighboring state CA, a different pattern was seen in the early pandemic where the lineage found early on in WA, CC256, only represented 20% of sequenced genomes at its most prevalent (1st-15th March) while CC4 (GISAID clade L) was the dominant variant, and was then replaced by CC258. Interestingly, a locally emerged variant CC10221 (GISAID clade G), probably from CC498, increased in abundance over time and then was likely exported to OR and NM (Supplementary Figure 2). A similar pattern was seen in WI where a local variant CC13301 increased in abundance over time and then appeared to spread to other states (NY, MI, MA and MN). In TX, multiple diverse CCs persisted in the population until mid-July.

In NY, a different pattern is seen with CC258 being persistently dominant. However, a more granular view of STs, not CCs, in New York shows a shifting epidemiology with ST258 declining and the rise of closely related single and double locus variants of ST258 reflecting local diversification (Supplementary Figure 3).

In MI, CC258 was the predominant strain until the summer when it gave way to a more diverse group of isolates. Similarly, in states like VA, CT, NM and LA mostly one predominant CC is seen over time, while in other states like UT, FL, OR, MA, MD and MN a diverse pattern of multiple CCs was noticed (Supplementary Figure 2).

The expansions and contractions in the temporal plots over time could be due to locally generated diversity (mutation) and/or introductions from other states or overseas. To better understand the source of ST diversity over time, we calculated indices reflecting effective circulating diversity as well as proportions of new STs in each state, and inferred domestic or global introductions and exportations based on previous observations in other locations or subsequent observations in other geographical locations (Figure 3B, Table 1 and Supplementary Figure 4). To infer introductions, we required that exactly the same ST was seen at least 10 days prior in some other geographical location. For exportations we required an ST to be seen first in the state in question at least 10 days prior to being seen anywhere else.

**Table 1.** Number of Genomes, Sequence Types, Simpson index, Hill Number, introductions and exportations for 16 US states.

| State | Genomes<br>(STs) | Simpson<br>Index ($^2H$) | Hill Number<br>($^2D$) | Non-<br>introductions | Introductions<br>(US) | Exportations |
| --- | --- | --- | --- | --- | --- | --- |
| WA | 3960<br>(1887) | 0.987 | 77 | 1817 | 44 (26) | 19 |
| TX | 2167<br>(1299) | 0.997 | 319 | 1258 | 31 (16) | 17 |
| CA | 1984<br>(1236) | 0.997 | 296 | 1173 | 35 (19) | 7 |
| NY | 1483<br>(825) | 0.960 | 25 | 766 | 25 (9) | 26 |
| MN | 1107<br>(522) | 0.988 | 81 | 470 | 29 (17) | 12 |
| WI | 954 (574) | 0.993 | 147 | 529 | 26 (15) | 8 |
| VA | 908 (543) | 0.994 | 165 | 511 | 18 (13) | 4 |
| LA | 850 (416) | 0.988 | 85 | 397 | 10 (10) | 1 |
| MI | 795 (416) | 0.889 | 9 | 384 | 16 (5) | 9 |
| FL | 750 (519) | 0.995 | 215 | 474 | 29 (18) | 6 |
| OR | 531 (343) | 0.995 | 190 | 320 | 19 (14) | 5 |
| UT | 350 (216) | 0.992 | 123 | 204 | 8 (4) | 2 |
| MA | 336 (170) | 0.940 | 17 | 144 | 17 (12) | 2 |
| MD | 196 (145) | 0.987 | 76 | 134 | 8 (4) | 2 |
| NM | 162 (109) | 0.987 | 80 | 103 | 3 (1) | 0 |
| CT | 154 (101) | 0.964 | 28 | 84 | 12 (8) | 0 |

The results of this analysis showed distinct patterns in different states with evidence supporting introductions usually outweighing evidence supporting exportations (Table 1). Interestingly, NY has the highest number of putative exportations (n=26), which was almost equal to the number of putative importations (n=25) potentially reflecting its role as a hub driving the initial pandemic. In most states there was a high amount of diversity that had no evidence of being introduced, which may signal problems with sampling, or may signal that local mutation is a strong force in generating diversity.

To understand the diversity within and between states, we calculated Hill numbers for all genomes from each state and over time in each state (Figure 4A, Table 1). Hill numbers are a diversity metric used widely in ecological studies that express effective diversity in units of sequence types, and they are less prone to biases introduced by incomplete or biased sampling (Alberdi and Gilbert 2019). Recognizing that our sample was not drawn from a systematically or evenly sampled dataset, we chose to use a Hill number metric (q=2) that emphasizes abundant taxa in estimating the effective diversity. Several other metrics such as the Shannon Index and a normalized richness index were highly dependent on the number of sampled genomes from each state. Hill numbers based on STs varied widely by state with TX showing the highest diversity and MI showing the lowest (Figure 3B and 4 and Table 1). Interestingly, there is a correlation (R^2^ = 0.1625) between effective diversity and when a state-wide mask mandate was imposed (Figure 4B).

**Figure 4.**
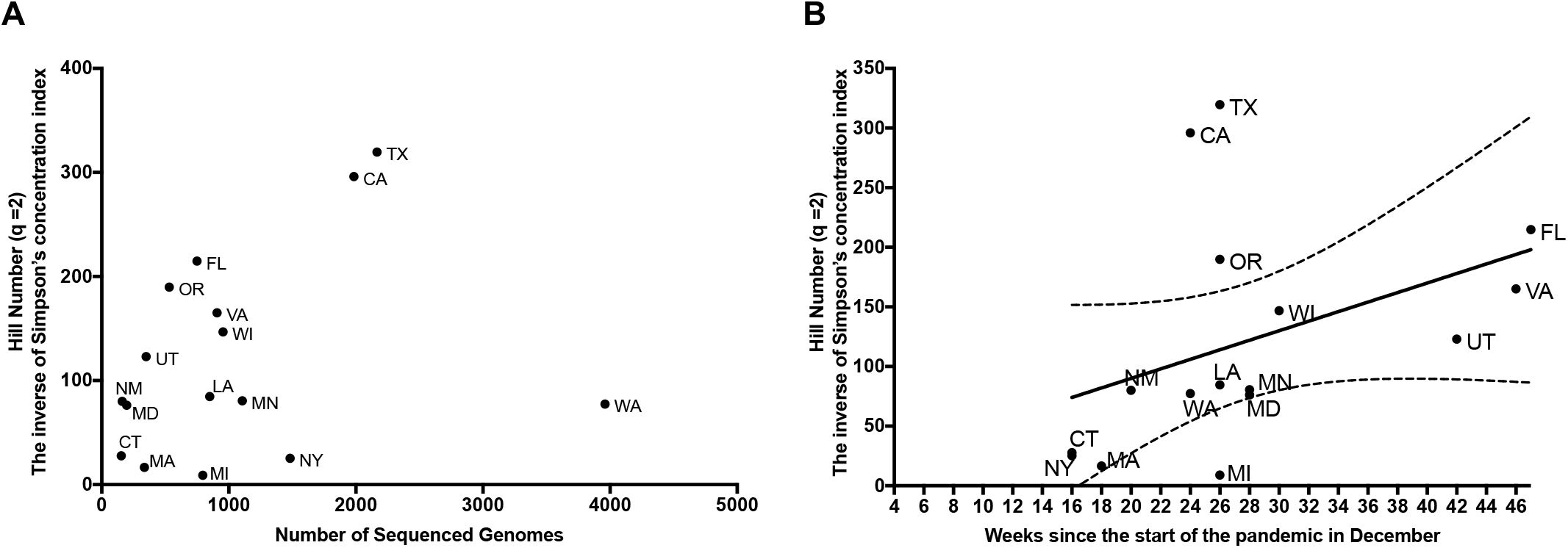
Effective Diversity of Sequence Types (STs) in 16 states. **A.** The Hill number equivalent (^2^D) of Simpson index (^2^H), is on the y-axis. Total number of genomes sequenced on the x-axis. **B.** Effective diversity (Hill number ^2^D) plotted against the week when state-wide mask mandate was imposed. Florida (FL) has no mask mandate so it was plotted at the end of the y-axis. The 16 different states are Washington (WA), California (CA), Wisconsin (WI), Texas (TX), New York (NY), Michigan (MI), Utah (UT), Virginia (VA), Florida (FL), Oregon (OR), Massachusetts (MA), New Mexico (NM), Maryland (MD), Connecticut (CT), Minnesota (MN) and Louisiana (LA).

Higher effective diversity may signal increased introduction of variants or increased local generation of new sequence types, which in turn may signal more open flow of virus into certain states or large circulating populations of virus able to mutate and diversify, respectively. To attempt to discriminate between these processes we calculated the effective diversity over time in each state and compared this to the proportion of novel variants that were determined to be introductions (Figure 3B and Supplementary Figure 4). In most states, initially high numbers of introductions were followed by a drop in the relative proportion of introductions as states began to impose restrictions in March. In some states the proportion of introductions also appears to increase over the summer as states eased regulations. Interestingly effective diversity also appeared to be correlated with peaks in the number of cases (Supplementary Figure 5) in several states, especially New York, but more data will be needed to be assessed to understand the connection between effective diversity and numbers of cases reported.

While our wgMLST approach is rapid and robust it has several limitations. Because a change in any allele creates a new ST our method may accumulate and count “unnecessary” STs that have been seen only once or may be due to a sequencing error. This is partially ameliorated by the use of the CC definition that allows some variability amongst the members of a group, and the use of only high-quality sequences. A large number of STs also may allow more granular approaches to tracking new lineages. Another limitation is the stability of the classification system, some virus genomes may be reassigned to new CCs as clones expand epidemiologically, but this may also reflect a dynamic strength as circulating viruses emerge and replace older lineages.

Perhaps most important limitation of our classification system is that it is limited by the quality and extent of the database. This is also reflected in the major limitation associated with the epidemiological and diversity inferences reported here. Uneven or biased sampling could lead to both inaccurate statements of the direction or origin of import/export events, and the source and quantification of diversity. The use of diversity statistics that emphasize more predominant variants and address sampling bias such as Hill numbers may help ameliorate this problem, but it seems clear that well-designed sampling strategies are needed to confidently understand ecological dynamics for SARS-CoV-2.

## Conclusion

The genomic epidemiology of the 69,686 SARS-CoV-2 isolates studied here show that 154 CCs have circulated globally and that more than half of these have been dynamically spreading through the US population with waves of changing diversity. Our tool (GNUVID) allows for fast sequence typing and clustering of whole genome sequences in a rapidly changing pandemic. As illustrated above, this can be used to temporally track emerging clones, identify the likely origin of viruses, and understand circulating diversity.

## Materials and Methods

All SARS-CoV-2 genomes (n=110,953) that were complete and have high coverage were downloaded from GISAID (Shu and McCauley 2017) on October 20^th^ 2020. Our wgMLST scheme was composed of all ten ORFs in the SARS-CoV-2 genome (Wu, et al. 2020). Genomes had to be at least 29,000 bp in length and have fewer than 1% “N”s. The ten ORFs were identified in the genomes using blastn (Altschul, et al. 1990) and any genome that had any ambiguity or degenerate bases (any base other than A,T,G and C) in the ten open reading frames (ORF) was excluded. The remaining 69,686 genomes (Supplementary table 1) were fed to the GNUVID tool in a time order queue (first-collected to last-collected), which assigned an ST profile to each genome. The identified STs by GNUVID were fed into the PHYLOViZ tool (Nascimento, et al. 2017) to identify CCs at the double locus variant (DLV) level using the goeBURST MST (Feil, et al. 2004; Francisco, et al. 2009). CCs were mapped back to the STs using a custom script. Pie charts were plotted using a custom script. The sci-kit learn implementation of Random Forest was then used to train a model. The model was trained using 53,565 SARS-CoV-2 sequences from GISAID representing the 154 CCs. Briefly, the 53,565 genomes were aligned to MN908947.3(Wu, et al. 2020) to generate a multiple sequence alignment using MAFFT’s FFT-NS-2 algorithm(Katoh, et al. 2002) (options: --add --keeplength). The 5’ and 3’ untranslated regions were masked in the alignment file using a custom script. Variant positions were then called using snp-sites (Page, et al. 2016) (options: -o -v). The 15,136 variant positions (features) matrix of the 53,565 CC-labelled genomes were then one-hot encoded, in which each SNP is replaced with a binary vector, and were used to train a random forest classifier in Scikit-learn (Pedregosa, et al. 2011). The prediction capability of the model was evaluated according to four statistics (accuracy, precision, recall and F-score).

To show the relationship between our typing scheme and phylogeny, we used a Global phylogeny of SARS-CoV-2 sequences from GISAID (last accessed 2020-11-13). The tree uses 99,160 high quality genomes(Lanfear and Mansfield. 2020). The tree and the 10 most common CCs were visualized in iTOL (Letunic and Bork 2019). We assigned a pangolin lineage (Rambaut, et al. 2020) (https://github.com/hCoV-2019/pangolin) and GISAID clade to each genome of the 53,565 genomes using the metadata details available on GISAID. We then compared the composition of each CC and calculated the percentage of the predominant clade/lineage in each CC (Supplementary table 1).

A total of 107,414 genomes (Supplementary table 1), that were training examples or assigned CCs and have date of isolation, were then used to analyze the number of introductions and exportations. Putative introductions were defined as an exact ST that was isolated somewhere else at least 10 days before the first date of isolation in the state in question. Exportations were defined as STs that were first isolated in the state in question and then isolated subsequently somewhere else at least 10 days later.

To compare diversity between the states and in each state over time, we calculated the Simpson index (Simpson 1949). To measure effective diversity in units of STs, we then transformed Simpson index (^2^H) to a Hill number (^2^D), which is the multiplicative inverse of the Simpson index (Alberdi and Gilbert 2019). The dates of state-wide mask mandates were the dates when face covering was required in indoor public spaces and in outdoor public spaces when social distancing is not possible (Abbott 2020; Allen 2020; Angell 2020; Baker 2020; Cuomo 2020; Edwards 2020; Evers 2020; Hogan 2020; Inslee 2020; Kunkel 2020; Lamont 2020; Northam 2020; Saunders 2020; Walz 2020; Whitmer 2020). The state-wide mandate dates used for WA, CA, TX, WI, NY, MI, LA, FL, MN, NM, OR, MA, MD, VA, UT and CT are 6/26/20, 6/18/20, 7/3/20, 8/1/20, 4/17/20, 7/10/20, 7/11/20, no mandate, 7/25/20, 5/16/20, 7/13/20, 5/6/20, 7/31/20, 12/14/20, 11/9/20, and 4/17/20,respectively. The Hill number is described as the effective number of STs (or CCs) of equally abundant STs (or CCs) that are needed to give the same diversity (Hill 1973; Jost 2006). The plots for number of confirmed cases in the 16 states were obtained from publicly available data in the Johns Hopkins University dashboard (Dong, et al. 2020).

The GNUVID database will be updated regularly with new added high-quality genomes from GISAID (Shu and McCauley 2017). Commands used are in Supplementary Methods. All the scripts are available from the authors and https://github.com/ahmedmagds/GNUVID (Moustafa and Planet 2020a). GNUVID can be installed through Bioconda (Grüning, et al. 2018).

## Availability of data and material

The compressed database and the trained model from our quality controlled genomes are available from the corresponding author and available online for download (Moustafa and Planet 2020a). The compressed database will be updated regularly on https://github.com/ahmedmagds/GNUVID. Source code for GNUVID can be found in its most up-to-date version here, https://github.com/ahmedmagds/GNUVID, under the GNU General Public License. All scripts are available from the authors.

## Authors’ contributions

Conceptualization: AMM, PJP; Coding: AMM; Writing – Reviewing and Editing: AMM, PJP.

## Acknowledgements

We would like to thank Ms Lidiya Denu, Dr Michael Silverman at the Children’s Hospital of Philadelphia, Mr Apurva Narechania at the American Museum of Natural History, and Dr. Joshua Mell Chang at Drexel University for helpful comments and discussion. We would like to thank the Global Initiative on Sharing All Influenza Data (GISAID) and thousands of contributing laboratories for making the genomes publicly available. A full acknowledgements table is available at https://github.com/ahmedmagds/GNUVID. This work was supported by the National Institute of Allergy and Infectious Diseases at the National Institutes of Health (1R01AI137526-01 and 1R21AI144561-01A1 to A.M.M. and P.J.P. and R01NR015639 to P.J.P.) and the Cystic Fibrosis Foundation (PLANET19G0 to A.M.M. and P.J.P.).

## Additional files

**Additional file 1:** Supplementary Methods and Figures.

**Additional file 2:** Table S1. GNUVID Database Strains Report Table.

