## Supplementary Methods and Figures for "Emerging SARS-CoV-2 diversity revealed by rapid whole genome sequence typing"

##### Installation using Bioconda

*$conda create -n GNUVID -c bioconda gnuvid*
*$conda activate GNUVID*

##### GNUVID Predict Usage

1. **Input**
   Query whole genome FASTA file (it can have multiple genomes as separate FASTA records).
2. **Command line options**
   a. **Required:**
   query_fna	Query whole genome nucleotide FASTA to analyze (.fna)
   b. **Optional:**
   -h Show help message and exit.
   -o Output folder and results prefix.
   -i	Individual output file for each genome showing the allele sequence and GNU score for each gene allele.
   -f Force overwriting existing results folder (default: off)
   -q No screen output (default: off)
   -v Print version and exit
3. **Output**
   a. **Always**
   GNUVID_results_date_time.csv (comma separated file, specify different name using -o option).
      - Column 1: Query Sequence name
      - Column 2: GNUVID Database version (results will vary as more genomes are added to the DB)
      - Columns 3-12: The allele numbers for the 10 ORFs (If None, it means the allele was not seen in the database but has degenerate bases (N) so cannot be called novel)
      - Column 13: ST
      - Column 14: First Country where the ST was seen (only if exact)
      - Column 15: First Date when the ST was seen (only if exact)
      - Column 16: Last Country where the ST was seen (only if exact)
      - Column 17: Last Date when the ST was seen (only if exact)
      - Column 18: Clonal Complex (CC) assigned
      - Column 19: Probability of the assignment (if exact, it means this is an exact match to a previous genome in the database)

   b. **Optional with -i**
   Genome1.csv (This report should have 10 rows for the ORFs. It will be produced for each genome. It is valuable if there is interest to know more about each ORF allele and how many times it was seen globally (GNU score) and when it was first- and last- time seen)
      - Column 1: Query Gene name
      - Column 2: GNUVID Database version (results will vary as more genomes are added to the DB
      - Column 3: GNU score (number of exact matches in the database, GNU=0 novel allele never seen before)
      - Column 4: Query gene sequence length
      - Column 5: Gene sequence

- Column 6: Number of Ns and degenerate bases in the query gene sequence
- Column 7: Alelle number from the database (If None, it means the allele was not seen in the database but has degenerate bases (N) so cannot be called novel)
- Column 8: First date this allele was seen (NA if novel)
- Column 9: Last date this allele was seen (NA if novel)

4. **Command example**
*$GNUVID_Predict.py new_genomes.fasta*

#### Commands Used for creating the compressed database and figures.

1. **Filter the fasta sequences by length and quality and cut into ORFs.**
*$GNUVID_FASTA_divider.py -l 29000 -N 1.0 GISAID_FASTA_10202020 gisaid_hcov-19_2020_10_20.fasta*
*$for i in `cat genomes.list`;do blastn -task blastn -out ${i}_results.txt -query MN908947.3_cds.fna -subject $i.fna -evalue 0.000001 -outfmt '6 qseqid sseq sstart send pident qcovs'; done*
*$Extract_fasta_sequence_blast_report.py ORFs _10202020 blast_reports_10202020/*
*$GNUVID_database_customizer.py -p -i -l db_genomes_list.csv db_fastas ORFs _10202020/*
*$GNUVID.py -m db_fastas/ -l strains_date_order.txt -o GNUVID_db_10202020 -p GNUVID_10202020 -cc country_continent.csv MN908947.3_cds.fna CDS test_fasta/*
2. **The identified STs by GNUVID were fed into the PHYLOViZ tool to identify CCs at the double locus variant (DLV) level using the goeBURST MST. CCs were mapped back to STs, pie charts were plotted and summarized using these commands.**
*$ Clonal_complex_assigner.py -n 20 -r resolve.csv GNUVID_10202020_DB_isolates_report_CC_assigned.txt ST_Full_MST.txt GNUVID_10202020_DB_isolates_report.txt*
*$Metadata_piechart.py pie_charts_CCs CC_list.txt GNUVID_10202020_DB_isolates_report_CC_assigned.txt*
*$GNUVID_CCs_summary.py 2020-09-20 2020-10-05 defining_SNPs_10202020.txt GNUVID_10202020_DB_isolates_report.txt*
3. **Training the random forest classifier.**
*$mafft --thread 32 --add Multifasta.fna --keeplength MN908947.3.fna > Multifasta_aligned.fna*
*$snp-sites -v -o SNPs_masked.vcf Multifasta_aligned_masked.fna*
*$snp-sites -o SNPs_masked.aln Multifasta_aligned_masked.fna*
*$GNUVID_Training.py 0.3 SNPs_masked.aln SNPs_masked.vcf*
4. **Analyzing 107,414 GISAID high coverage genomes, predicting the number of introductions/exportations and temporally plotting ratios reflecting circulating diversity, novel STs and introduced STs.**
*$GNUVID_Predict.py gisaid_107414_isolates.fasta*
*$Extract_US_genomes.py WA TX CA WI NY MI MN LA UT VA FL OR MA NM MD CT GISAID_107414_genomes.txt*
*$chmod +x Temporal_plot_collector_curve.sh*
*$./Temporal_plot_collector_curve.sh*
5. **Calculating Hill Number $^2$D (Simpson Diversity) for each state.**
*$Alpha_diversity.py WI WI_reports_order.txt WI_GNUVID_reports/*

#### Supplementary Figures

**Supplementary Figure 1**

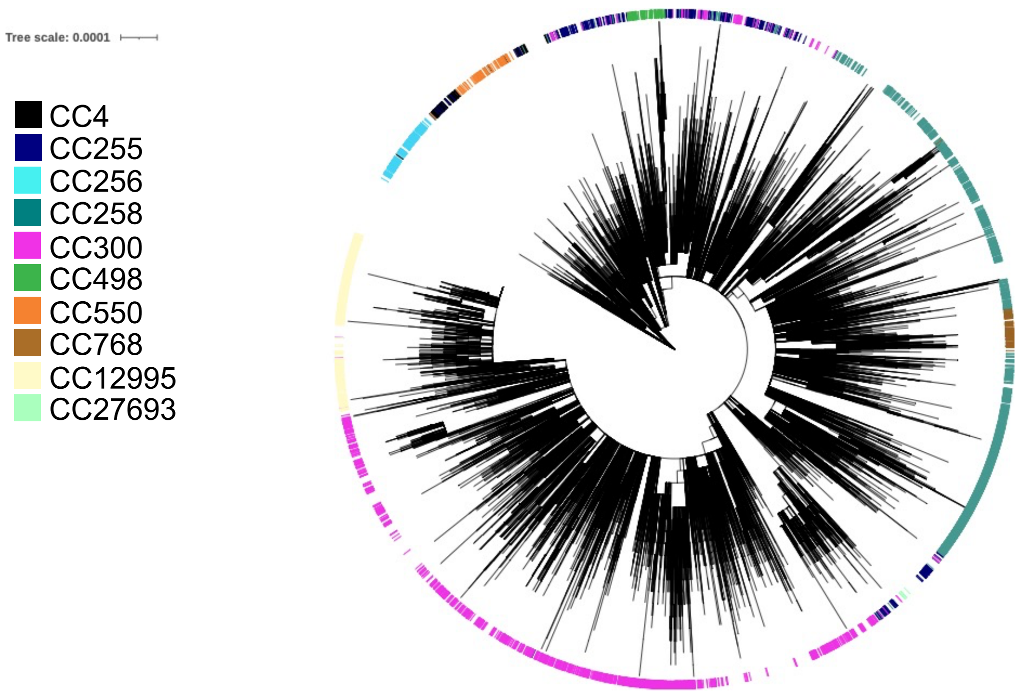

Supplementary Figure 1. Maximum likelihood global phylogeny of 99,160 SARS-CoV-2 genomes. The tree and 10 most common CCs data were visualized in iTOL. Members of the same CC usually grouped together in clades.

#### Supplementary Figure 2

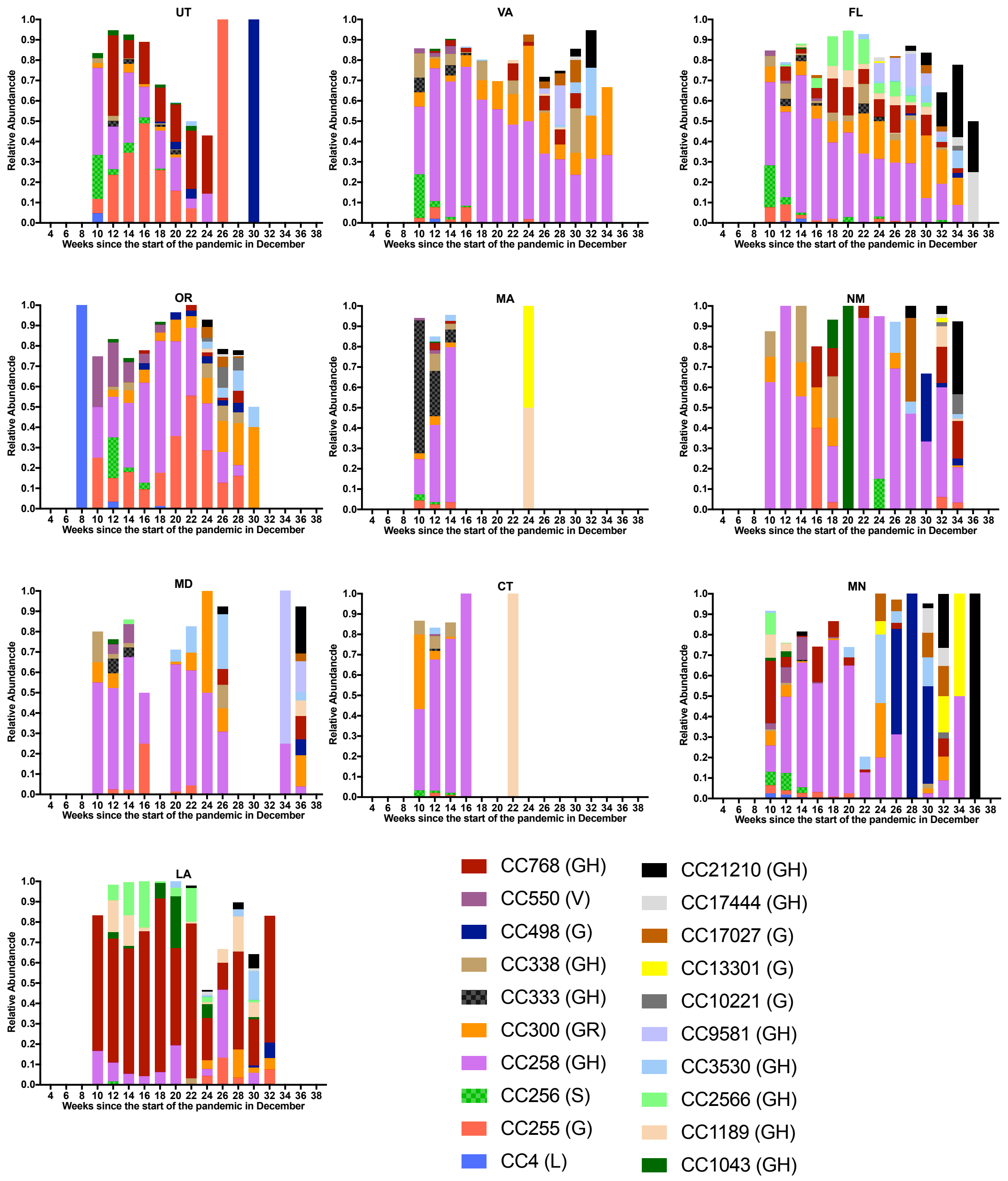

Supplementary Figure 2. Temporal Plots of circulating Clonal Complexes and corresponding GISAID clade in parentheses at 10 different states (Utah (UT), Virginia (VA), Florida (FL), Oregon (OR), Massachusetts (MA), New Mexico (NM), Maryland (MD), Connecticut (CT), Minnesota (MN) and Louisiana (LA)). The visualizations were limited to the 20 most common CCs.

**Supplementary Figure 3**

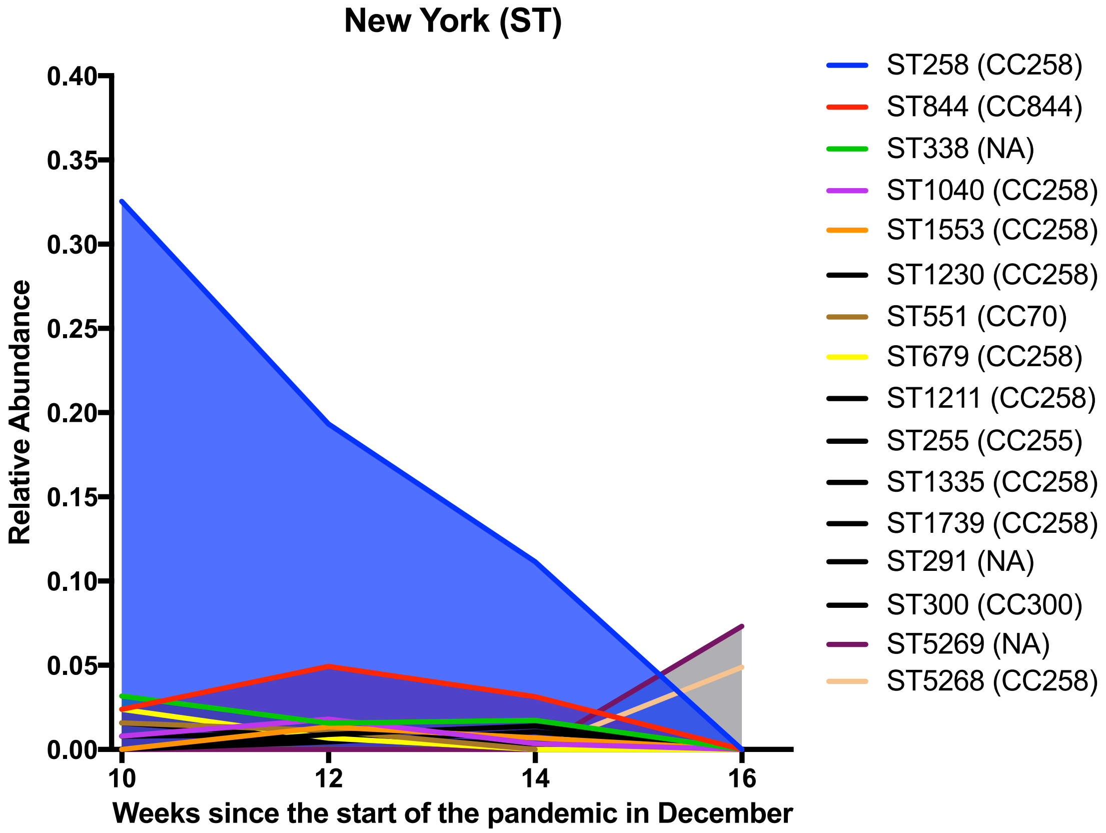

Supplementary Figure 3. Temporal Plot of circulating Sequence Types (STs) at New York (NY).

#### Supplementary Figure 4

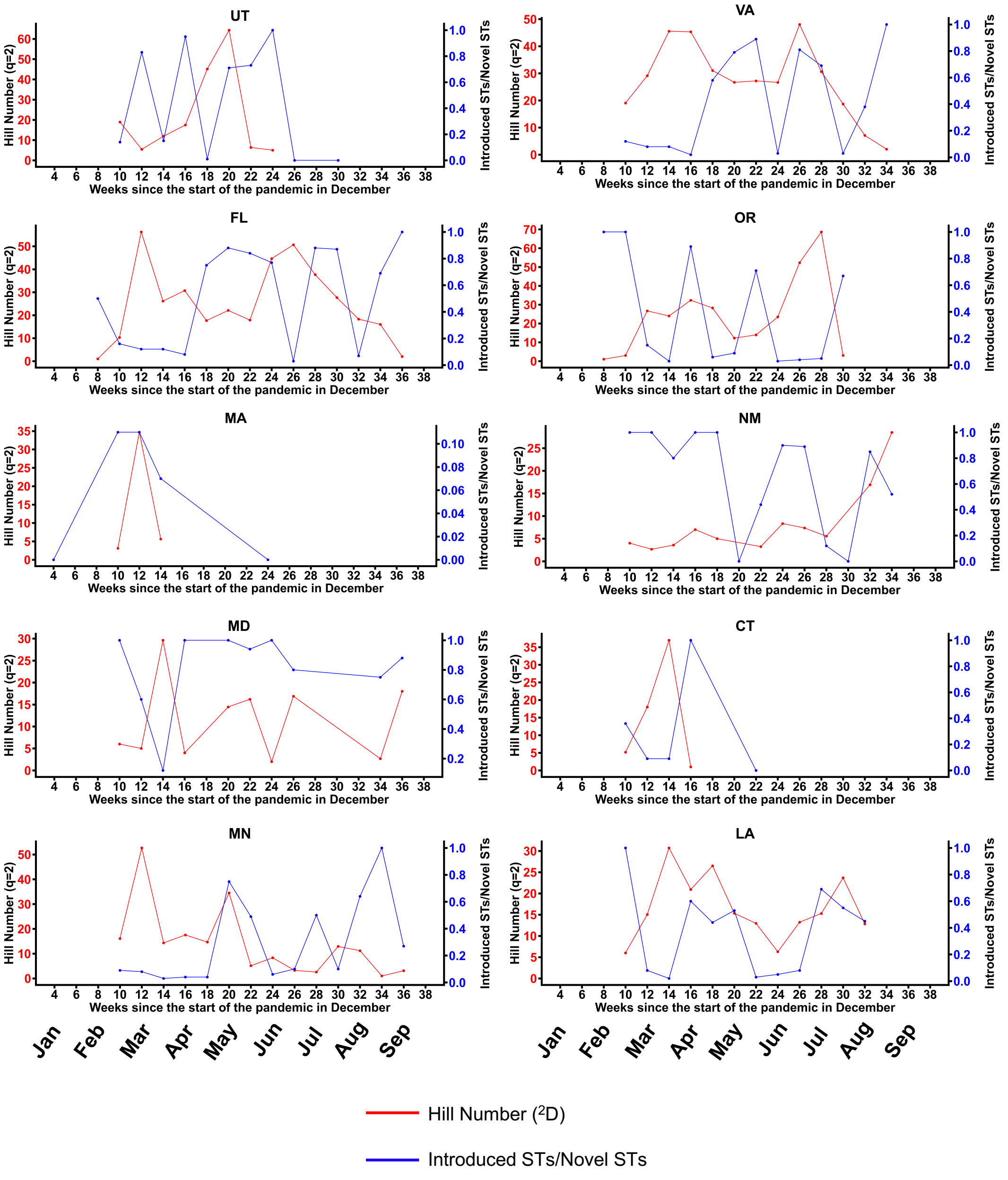

Supplementary Figure 4. Diversity of Sequence Types (STs) in 10 different states (Utah (UT), Virginia (VA), Florida (FL), Oregon (OR), Massachusetts (MA), New Mexico (NM), Maryland (MD), Connecticut (CT), Minnesota (MN) and Louisiana (LA)) over time are represented for each 2-week time period in the following ratios: (1.) Effective diversity (Hill number equivalent ($^2$D) of Simpson index ($^2$H)) (red) (2.) Number of STs putative introductions for each time period divided by the total number of STs not seen previously in a state (blue).

### Supplementary Figure 5

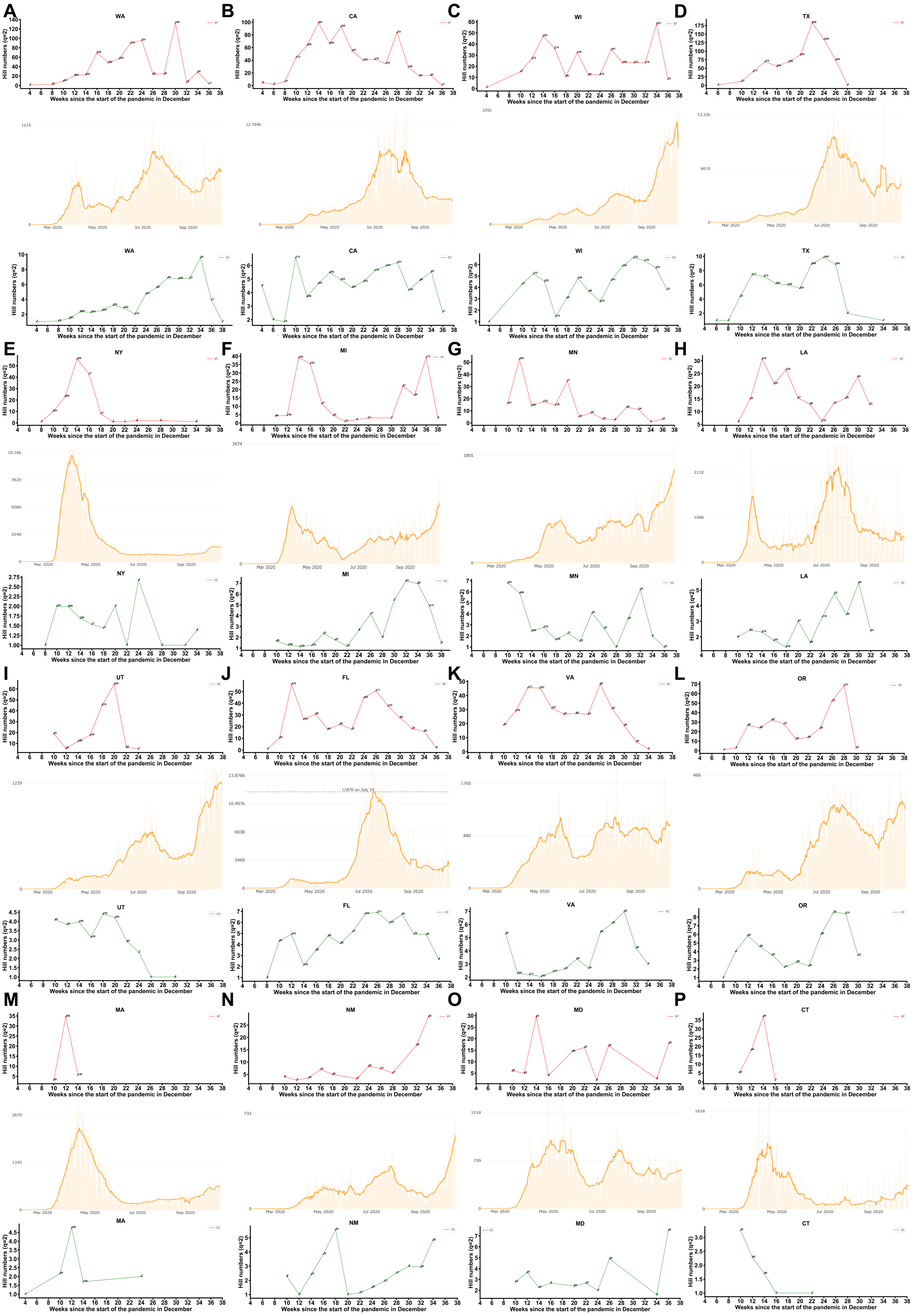

Supplementary Figure 5. Effective Diversity of Sequence Types (STs) and Clonal Complexes (CCs) in the 16 states over time and the corresponding number of confirmed cases based on data from Johns Hopkins University dashboard. The Simpson diversity for STs and CCs are represented for each 2-week time period in red and green, respectively. The plots for number of confirmed cases in each of the 16 states are shown. The 16 different states are Washington (WA), California (CA), Wisconsin (WI), Texas (TX), New York (NY), Michigan (MI), Utah (UT), Virginia (VA), Florida (FL), Oregon (OR), Massachusetts (MA), New Mexico (NM), Maryland (MD), Connecticut (CT), Minnesota (MN) and Louisiana (LA).
